# Establishing correlation of axial and vertical force of cardiac artificial tissue using a novel micromachined sensor

**DOI:** 10.1101/2020.09.16.300665

**Authors:** Angelo Gaitas, Francesca Stillitano, Irene Turnbull

## Abstract

Cardiomyocytes iPSC (iPSC-CMs) have great potential for cell therapy, drug assessment, and for understanding the pathophysiology and genetic underpinnings of cardiac diseases. Contraction forces are one of the most important characteristics of cardiac function and are predictors of healthy and diseased states. Cantilever techniques, such as atomic force microscopy, measure the vertical force of a single cell, while systems designed to more closely resemble the physical heart function, such as cardiac tissue on posts, measure the axial force. One important question is how do these two force measurements correlate? By establishing a correlation of the axial and vertical force we will be one step closer in being able to use single cell iPSC instead of more elaborate human engineered tissue or animal heart tissue as models. A novel micromachined sensor for measuring force contractions of artificial tissue has been developed. Using this novel sensor a correlation between axial force and vertical force is experimentally established. This finding supports the use of vertical measurements as an alternative to tissue measurements.

## 1. INTRODUCTION

Human induced pluripotent stem cells (iPSC) are pluripotent stem cells derived from adult cells (e.g. adult human skin cells) and converted into other cell types, such as cardiomyocytes (CM) [1]. Cardiomyocytes iPSC (iPSC-CMs) have great potential for cell therapy[2, 3], drug assessment[4], and for understanding the pathophysiology and genetic underpinnings of cardiac diseases[5]. iPSC-CMs are emerging as a promising platform for drug evaluation[6] and hold significant potential for use in predicting cardiotoxicity[7, 8] and may find uses in personalized medicine [XX]. Of particular importance is the use of iPSC-CM as a human cellular model for predicting drug arrhythmogenicity [9]. Nearly one third of all drugs have cardiovascular side effects [10, 11].

Contraction forces are one of the most important characteristics of cardiac function and are predictors of healthy and diseased states[7, 12]. Several techniques have been used to measure the force of contraction from cardiomyocytes [7, 13-22]. However, most of these techniques are not useful in characterizing iPSC-CMs due to their irregular shape and smaller size, also many of these techniques are unable to directly measure the force. An answer to this problem is the use of microcantilevers, such as atomic force microscopy (AFM) cantilevers [23, 24] and micro-cantilevers with embedded sensors [25, 26], that have already been used for cardiomyocyte contractile force measurements. These devices are like micron size diving boards that are brought in contact with single cells. Either using a laser light that is reflected off the cantilever, or by using an embedded sensor, the movement of the cantilever that corresponds to the cellular contractions is tracked. By characterizing the properties of the microcantilever it is a simple task to derive the force.

The cantilever techniques measure the vertical force of a single cell, while systems designed to more closely resemble the physical heart function, such as cardiac tissue on posts[27, 28], measure the axial force (Fig. 1 (a)). One important question is how do these two force measurements correlate? By establishing a correlation of the axial and vertical force we will be one step closer in being able to use single cell iPSC instead of more elaborate human engineered tissue or animal heart tissue as models. This would enable the use of single iPSC for cardiotoxicity and personalized medicine. In this work, we attempt for the first time to establish such a correlation through measurements of human engineered tissue axial force using an established method [29] and vertical force using a micromachined cantilever with an embedded strain sensor. These measurements are not possible with a conventional AFM due to the AFM’s travel range limitations and large size of tissue. While still preliminary, our results point to the existence of such a correlation.

**Fig. 1.**
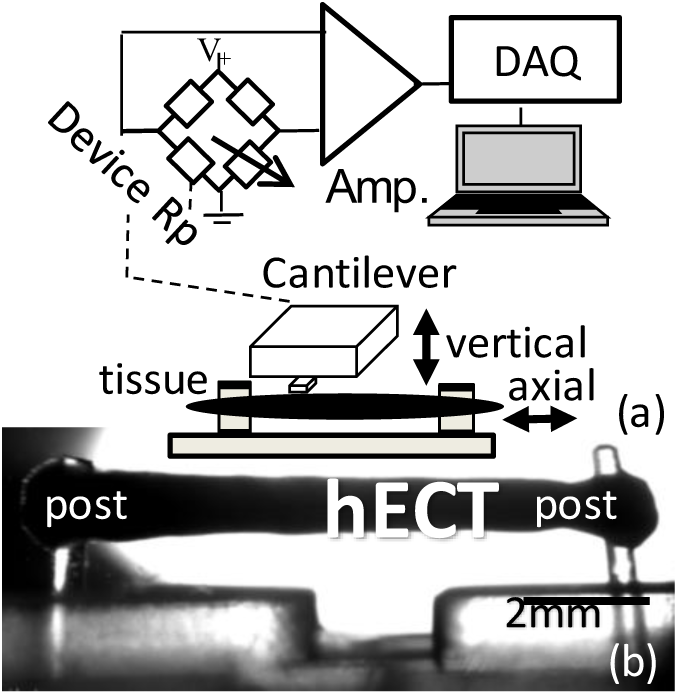
a) Schematic diagram of the experimental setup. A Wheatstone bridge circuit with one of the resistors being the strain gauge (force) sensor of the microcantilever, the voltage signal is amplified and fed into a DAQ card. On the bottom diagram a cantilever is shown in contact with the tissue. The vertical and axial force directions are schematically shown. b) Picture of hECT on posts.

## 2. MATERIAL AND METHODS

### 2.1 Micromachined force sensors and interface circuitry

A specialized micromachined cantilever force sensor was fabricated for these measurements similar to the ones published previously (***Fig. 1***) to measure contractile forces [25, 26]. The sensor operates like an electronic finger, detecting contact with a cell and cellular or tissue contractions. The body of the cantilever sensor is made of polyimide and includes a thin film strain gauge made from Cr/Au with 1 nm/10 nm thickness that is sandwiched between two layers of polyimide, 1.5 μm and 0.5 μm, respectively. The sensor is 40 μm wide, 150 μm long, and approximately 2 μm thick. The fabrication process requires four masks. The detailed fabrication process flow is shown in **Figure 2**. In brief, a thermal oxide mask is deposited and patterned (1). The bottom layer of polyimide is spin-coated and patterned with reactive-ion etching (RIE) (2). A thin Cr/Au (1nm/10nm) metal line is sputtered for the resistor and a thicker Cr/Au (1 nm/ 100nm) is deposited on the pad area for electrical bonding (3). The top layer of polyimide is spin-coated and patterned with RIE (4). The cantilever structure is formed by a backside deep reactive-ion etching (DRIE) process (4). The oxide acts as an etch stop. Finally, the thermal oxide layer is removed in buffered oxide etch (BHF) (6). Al is deposited on the backside of the wafer for refection. The resulting device is shown in Fig. X(b). The inset of Fig. 2(b) shows the device chip which includes two sensor cantilevers one of which is broken off. The device chip is wired and passivated using PDMS. The sensor’s spring constant, k, is ∼0.059 N/m and its nominal resistance is between 160 and 170 Ω.

**Fig 2.**
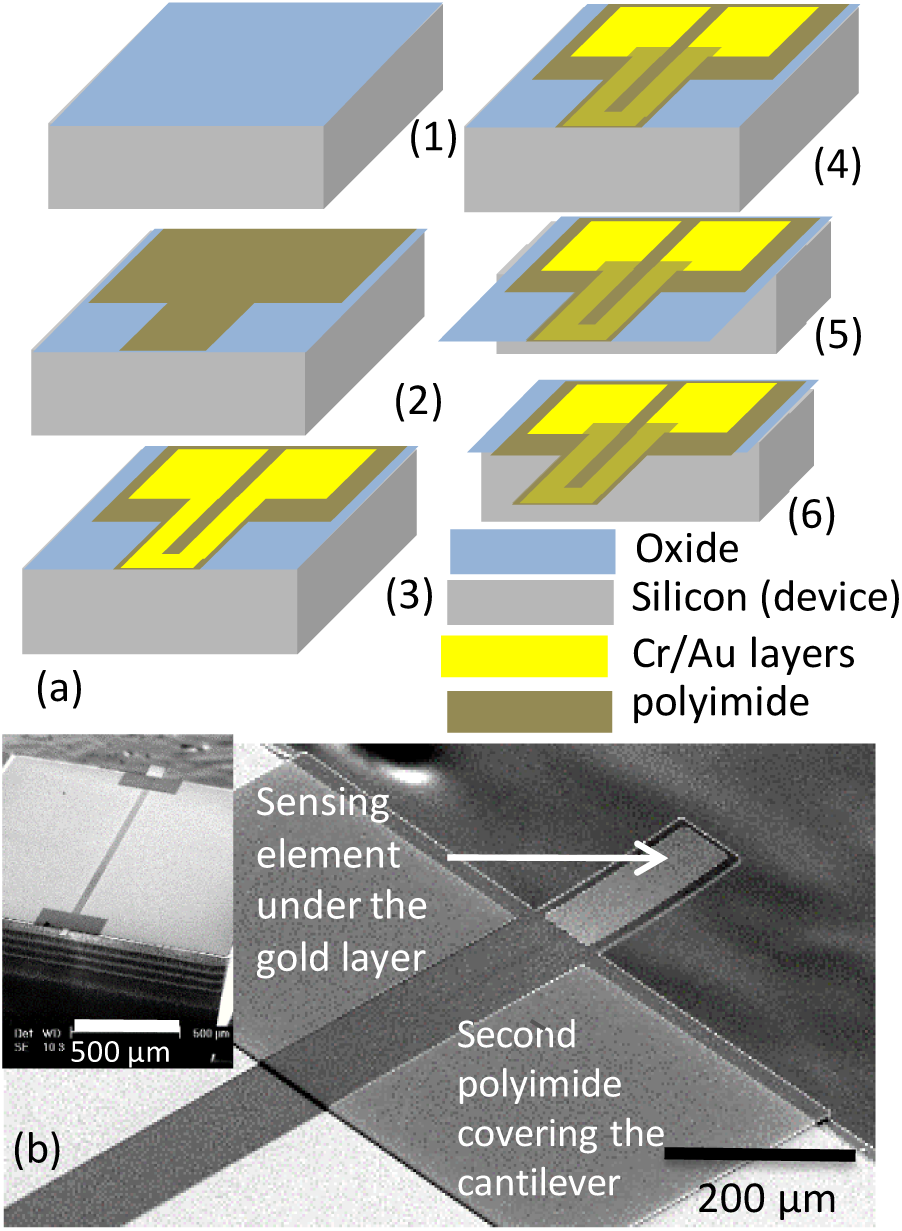
a) Fabrication steps to produce the micro-cantilever force sensor. b) Scanning electron micrograph of the cantilever device (Inset: die with two devices one of which is broken off).

The movement of the cantilever is controlled with a motorized micromanipulator with 25 × 25 × 25 mm^3^ range and 0.1 μm minimal increment step (KT-LS28-MV, Zaber Technologies). Lab-VIEW software was used to control the movement and record the output of the sensor. The electrical signal from the cantilever was measured using a Wheatstone bridge circuit connected to a low noise amplifier, and a National Instruments digital acquisition (DAQ) card to record the data into a computer (Fig. 1 (a))). The cantilever was calibrated by bringing it in contact with a hard surface such as a glass slide and recording the change in voltage from the bridge circuit output with distance travelled in the Z-axis. Multiplying the spring constant of the microcantilever with the distance travelled the force can be estimated.

### 2.2 Human engineered cardiac tissue and optical posts tracking

The induced pluripotent stem cell (iPSC) line (SKiPS-31.3) was derived from adult dermal fibroblasts, which were obtained with informed consent from a healthy adult volunteer [30]. This research project involved only secondary analysis of already collected biological samples without personal identifiers. iPSCs were grown in StemFlex media on tissue culture 6-well plates coated with hESC-qualified Matrigel in 5%CO^2^ at 37°C. iPSCs were differentiated into CMs by following a monolayer-based differentiation protocol [29]Briefly, when iPSCs reached ∼80% confluence, StemFlex media was replaced with basal medium (RPMI 1640 media plus 2% B27 supplement minus Insulin), and 10 μM CHIR99021 for 24 hours. On day 1, the media was replaced with basal medium without CHIR99021 for two days. On days 3 and 4, media was replaced with basal media with 5 μM IWR-1. On day 5, the media was replaced with basal media. On day 7, basal medial was replaced with RPMI 1640 media plus 2% B27 supplement. Afterward the media was exchanged every 2 days. Functional iPS-CMs appear in culture between days 7 and 10 post-CHIR treatment.

For hECT fabrication, hiPSC derived cardiomyocytes (hCM) were collected after 24 days of differentiation, by enzymatic digestion of the hCM monolayer (using trypsin 0.25%); the hCM cell pellet was resuspended in a solution of type-I collagen and Matrigel, 100ul of this cell-matrix mix was layered into the well of a single-tissue bioreactor [29] with each resulting hECT containing approximately 1.5 x10^6^ cells. The bioreactor was placed in a 60mm dish and after 2 hour incubation for polymerization, the bioreactor was covered with culture medium (RPMI 1640 media plus 2% B27 supplement), and maintained at 5% CO2 and 37°C with half-media exchanges every other day.

The single-tissue bioreactor is fabricated of polydimethylsiloxane (PDMS) with flexible endposts (Young’s modulus 1.33MPA), the posts serve as anchors during formation of the hECT, and act as integrated force sensors that deflect during hECT beating [29, 31]. The average Young’s modulus of the posts was 1.33 MPa from calibration tests using a high-sensitivity force transducer (Scientific Instruments, Heidelberg, Germany). The tissues typically begin to beat spontaneously within 72 hours, and deflection of the end posts with each contraction can be visualized with low magnification microscope by day 6. The axial force of the engineered tissues is measured using an established method to optically track end posts for force calculation described in [29](***Fig. 1 (b)***). The contractile function of each hECT was measured in a laminar flow hood in a custom setup that allows optical tracking of PDMS post deflection versus time using custom LabVIEW software to acquire real-time data sampling. The data is then analyzed with a custom MATLAB script to calculate twitch parameters including developed force (DF; calculated from the post deflection with each twitch), and developed stress (DS; DF divided by hECT cross sectional area), as previously described [29]. Recordings were first performed without electrical stimulation to obtain data on spontaneous beating frequency. Then the hECTs were electrically paced by field stimulation (12-V biphasic pulse with 5-ms duration) at increasing frequencies, using a programmable Grass S88X stimulator (Astro-Med, West Warwick, RI).

### 2.3 Experimental set-up

The microcantilever was positioned in close proximity to the hECT and brought it in contact under optical observation with the tissue using the motorized stage (***Fig. 3(a) & (b)***). Due to the large range of motion afforded by the system it is relatively simple to achieve contact with the sample. While moving the microcantilever closer to the sample the voltage signal coming out of the bridge circuit is monitored. A sudden increase in the voltage is indicative of contact. Once in contact, contractions start being recorded as shown in *Fig. 3(c)*. The voltage changes of the microcantilever are monitored and plotted using LabVIEW (a visual programming language) (**Fig. 3(c)**). The vertical force measured by the microcantilever to the axial force measurements are compared to determine the relationship between these two forces.

**Fig 3.**
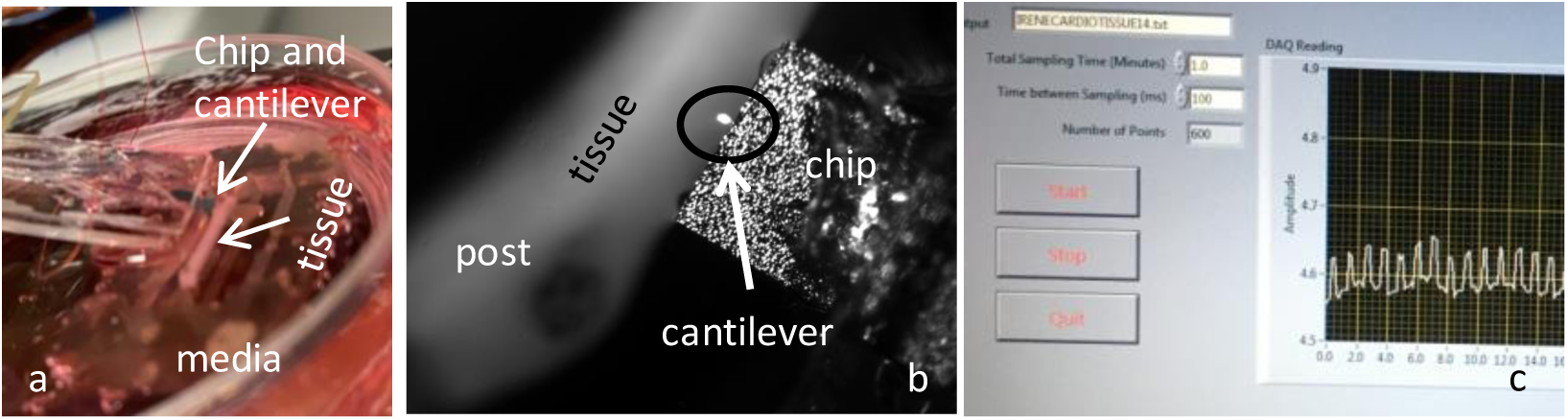
a) Picture of microcantilever immersed in media and in contact with hECT. b) Picture of microcantilever approaching the hECT. c) Screenshot of real time LabView recording of microcantilever voltage changes as the tissue in contact contracts.

## 3. RESULTS AND DISCUSSION

The tissue was paced between 0.2 and 1 Hz and the microcantilever was able to accurately measure tissue contraction frequency changes as shown in ***Fig. 5***. Similar to what it is observed using the post, the magnitude of the force measured by the microcantilever drops with increasing frequency. The relationship is linear at small frequency values below 1 Hz *(N= 6, error bars = STD)* (***Fig. 5 (f))***. Linear regression was used to examine the association between force and frequency: Force per unit area = −0.31 frequency + 0.36 (R^2^ = 0.99). The results demonstrate that the microcantilever can measure frequency in response to pacing hECT and force.

**Fig. 5.**
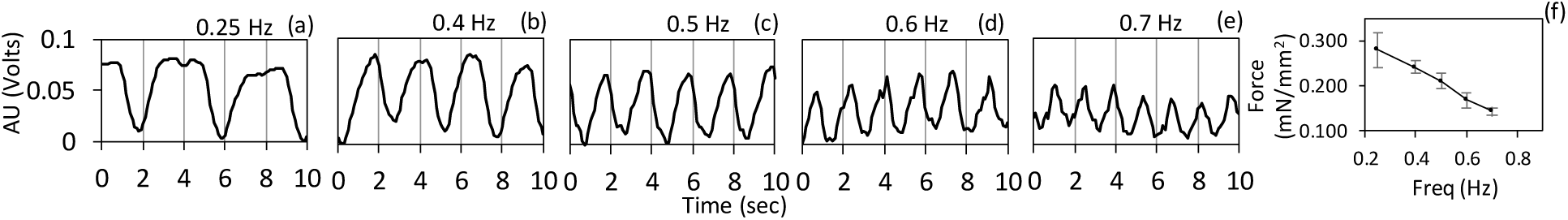
(a-e) In the Y-axis the cantilever reading in volts from 0-0.1V. This is the reading from the Wheatstone bridge circuit that was amplified x1000. In the X-axis the time is plotted in seconds. (f) Plot of force per unit area with frequency. At this small frequency range the change is highly linear. As the frequency increases the amplitude of the contraction and the force decrease. (N= 6, error bars = STD).

We examined the association between the force measure from the post (axial force - AF) in the y-axis and the cantilever force measured in the x-axis (vertical force – VF) with linear regression (***Fig. 6***). A significant regression equation was found (AF (1, 3)= 169.5, p < 0.001) with R^2^ of 0.98. The axial force AF is equal to 0.071 + 0.248 VF. Thus the vertical force is a significant predictor of the axial force. Our experimental findings suggest that the axial and the vertical force are strongly correlated. These results are significant because for the first time they establish a relationship between axial and vertical force measured by the MEMS cantilever.

**Fig. 6.**
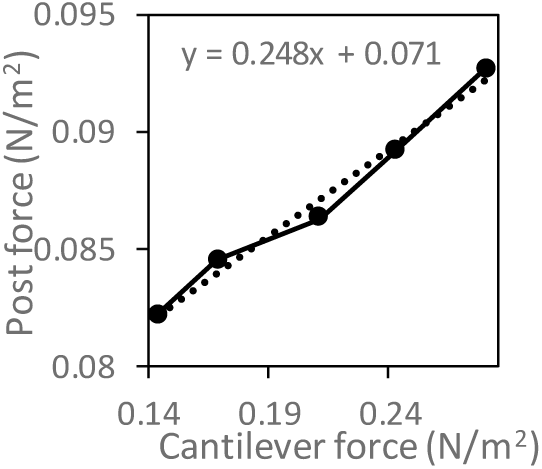
Plot of force from the posts in the y-axis vs. the cantilever force measured in the x-axis. Inset include linear regression model.

## 4. CONCLUSIONS

In this work the relationship between the vertical force measured by a microcantilever device and the axial force in human engineered cardiac tissue (hECT) measured using established techniques has been shown. These results support the use of single cell vertical force measurements as an alternative to axial force measurements.

## Abbreviations

SPM: scanning probe microscopy
AFM: atomic force microscopy
CM: cardiomyocyte
iPSC: induced pluripotent stem cells
MEMS: micro-electromechanical systems
hECT: human engineered cardiac tissue

## ACKNOWLEDGEMENTS

This work was supported by the National Institutes of Health (U24 DK112331-03S1 and K01HL133424).

